# Adaptor Template Oligo-Mediated Sequencing (ATOM-Seq): A versatile and ultra-sensitive UMI-based NGS library preparation technology, for use with cfDNA and cfRNA

**DOI:** 10.1101/2020.07.10.152405

**Authors:** Thomas L. Dunwell, Simon C. Dailey, Jihang Yu, Philipp W. Becker, Sarah Scaife, Susan D. Richman, Henry M. Wood, Hayley Slaney, Daniel Bottomley, Xiangsheng Yang, Hui Xiao, Anine L. Ottestad, Sissel G. F. Wahl, Bjørn H. Grønberg, Hong Yan Dai, Guoliang Fu

## Abstract

Liquid biopsy testing utilising Next Generation Sequencing (NGS) is rapidly moving towards clinical adoption for personalised oncology. However, before NGS can fulfil its potential any novel testing approach must identify ways of reducing errors, allowing for separating true low-frequency mutations from procedural artefacts, and be designed to improve upon current technologies while also avoiding their limitations. Popular NGS technologies typically utilise two approaches; PCR and ligation, which have known limitations and seem to have reached a development plateau with only small, stepwise improvements being made. To maximise the ultimate utility of liquid biopsy testing we have developed a highly versatile approach to NGS: Adaptor Template Oligo Mediated Sequencing (ATOM-Seq). ATOM-Seq’s strengths and versatility avoid the major limitations of both PCR- and ligation-based approaches. This technology is ligation free, simple, efficient, flexible, and streamlined, and offers novel advantages that make it perfectly suited for use on highly challenging clinical material. Using reference materials, we demonstrate detection of known SNVs at allele frequencies of 0.1% using as little as 20 ng of DNA, as well as the ability to detect fusions from RNA. We illustrate ATOM-Seq’s suitability for clinical testing by showing high concordance rates between paired cfDNA and FFPE clinical samples.

## Introduction

The ever-expanding use of next generation sequencing (NGS) has been instrumental in exploring and understanding the changes which occur during cancer development. This has revealed expansive patterns of single-nucleotide variants (SNVs), insertions, deletions, copy number variations (CNVs) and gene fusions as frequently occurring events across one or more cancer types, an approach epitomised by studies such as The Cancer Genome Atlas [1]. The understanding this has given us about cancer development has motivated the development of both PCR- and NGS-based technologies for use in the detection of these cancer-associated changes. Historically these technologies and associated tests have focused on identifying mutations through testing of tumour biopsies, and with these materials previous technologies have performed well. Testing of primary tumours is relatively simple, but obtaining this material can be extremely invasive. In an attempt to improve patient outcomes, attention has therefore turned towards using minimally invasive liquid biopsies. This is a highly challenging approach which is motivated by the fact that cancer-related biomarkers can be detected in these easily obtained blood samples without any prior knowledge of the location of a tumour and potentially at a far earlier stage of disease progression.

There are wide ranging potential benefits to liquid biopsy testing, as these samples can contain a combination of cell free DNA and (to a lesser extent) cell free RNA (cfDNA/cfRNA) derived from primary tumours; clinical testing of these molecules allows for the indirect detection and genotyping of tumours (including SNV/CNV/fusions). The ability to interrogate these samples holds great promise for early diagnosis and for disease monitoring in relation to determining the success of targeted therapies and detecting the emergence of drug resistance clones, as well as in assessing disease recurrence and minimal residual disease monitoring. Unfortunately, the majority of the material in a liquid biopsy will have derived from non-cancerous cells. As a result, DNA/RNA from a tumour may only be present at a very low proportion (0.01%) of the total quantity of nucleic acids in a sample. This poses a significant challenge, as the apparent allele frequency of the mutation may fall below the background error rate for a given sequencing technology. Successfully demonstrating clinical validity and clinical utility of testing liquid biopsy-derived material using NGS will therefore depend on the availability of technologies that are able to detect cancer-associated changes with very high sensitivity and specificity [2].

When considering the NGS technologies currently available, these can be broadly categorised as ‘whole genome’ or ‘targeted’ approaches. The depth necessary to identify rare and ultra-rare variants (<0.5-1%) makes whole genome sequencing costly and impractical, and as such the majority of adopted approaches have focused on targeted technologies for cancer-associated mutation testing. The detection of low frequency mutations by these target enrichment processes are hampered by a low level of background errors, which are present in all workflows [3-7]. To combat this, technologies have been adapted to incorporate unique molecular identifiers (UMI). These are short regions of nucleic acids that are added to original DNA fragments, or copies thereof, in the first step(s) of an enrichment protocol. After sequencing, the UMIs are used to identify PCR products known to come from the same original molecule, as each will share the same UMI. These are then de-multiplexed and as a result amplification errors can be corrected, resulting in the identification of a series of consensus reads which represent the original starting material. By taking this approach, many protocol-based errors (such as mis-incorporation of nucleotides by polymerases) can be significantly reduced.

These UMI-containing enrichment technologies can broadly be separated between 1) those approaches which use PCR to directly copy and amplify the original DNA [8], and 2) those based on ligation of adaptors onto the ends of double-strand DNA. The latter approach can be further subdivided into those which either undergo subsequent probe based capture [9,10] or PCR-based enrichment using ligation products as a template [11-13]. All of these methods have been utilised for testing clinical samples for ultra-rare variants, especially when using liquid biopsy samples. cfDNA in liquid biopsy samples is enriched in short, single-strand, and damaged molecules [14-16] and this results in issues for these technologies. The reliance of purely PCR-based approaches on a predetermined pair of opposing primers means these approaches are ill-suited to identifying large indels and translocations or to heavily fragmented input material. These approaches also cannot detect unknown gene fusions. Ligation-based approaches used in targeted enrichment protocols are generally double-strand DNA specific, meaning they are incapable of working with single-strand DNA and also struggle with damaged DNA. Approaches have been designed to be more efficient with ssDNA [17-19]. The ligation-hybrid capture approaches struggle with capturing short DNA fragments and are also very long and complex protocols. The inherent limitations of the aforementioned technologies in processing liquid biopsy samples means they will inevitably fail to maximise their sensitivity in detecting cancer-associated changes obtained from such clinical samples. As such there is a need for the development of new approaches which are specifically designed for liquid biopsy testing and are focused on overcoming these limitations.

With both the limitations of current technologies and biological context of cell free nucleic acids in mind, we re-imagined the process of library preparation and developed a novel technology designed specifically for use with challenging clinical material such as cfDNA/cfRNA obtained from liquid biopsies: **<u>A</u>**daptor **<u>T</u>**emplate **<u>O</u>**ligo **<u>M</u>**ediated **<u>Seq</u>**uencing - ATOM-Seq.

In this paper, we describe ATOM-Seq’s technological approach for NGS library preparation and detail its highly efficient, robust and rapid protocol, demonstrating suitability for both targeted and whole genome library preparation methods, using RNA or DNA nucleic acids as starting material. We demonstrate the technology’s sensitivity using reference material for detecting rare variants down to 0.1% allele frequency using as little as 20ng of fragmented DNA and show that it has sensitivities and specificities >97%. With clinical samples we show excellent performance on colorectal cancer formalin-fixed paraffin-embedded (FFPE) samples, lung cancer liquid biopsy samples and FFPE samples when compared with previous technologies, and with paired lung cancer FFPE and liquid biopsy samples we demonstrate concordance rates >83% for de novo mutation detection. With RNA we demonstrate the technology’s suitability for detecting gene fusions, and with total nucleic acid samples (combined cfRNA and cfDNA) we demonstrate an ability to capture and enrich for both cfRNA and cfDNA molecules in a single reaction.

## Results

### Adaptor Template Oligo Mediated Sequencing

The main goal of this project was to design a new approach for liquid biopsy material testing. We successfully developed a new NGS library preparation technology we named **<u>A</u>**daptor **<u>T</u>**emplate **<u>O</u>**ligo **<u>M</u>**ediated **<u>Seq</u>**uencing - ATOM-Seq. This technology was designed to be compatible with any suitable starting material including single- and double-strand DNA, cfDNA, enzymatically fragmented or sonicated gDNA or FFPE DNA, and single- and double-strand cDNA produced from FFPE and cell free RNA.

The underpinning conceptual idea of ATOM-Seq was to use the original sample DNA as a primer and extend its 3’ end, thereby allowing ‘capturing’ of every molecule in the starting material. This was accomplished by designing an approach where the starting material is combined with a synthetic oligo, termed an Adaptor Template Oligo (ATO). The 3’ ends of the starting material anneal to a single-strand random sequence contained on the 5’ end of the ATO. Through the addition of a suitable polymerase the starting material then functions as a primer and is extended using the ATO as a template (Figure 1A). This 3’ extension reaction copies the DNA sequence from the ATO onto the 3’ end of the starting material, ‘capturing’ the DNA ends. During this reaction two separate functional components are created on each 3’ end. The first is a random sequence which functions as a true Unique Molecular Identifier (UMI), allowing for bioinformatic error correction after sequencing. The second is a universal primer site which is used during downstream amplification reactions. The results of these reactions, which we have termed ATO-Reactions, are conceptually similar to those generated with traditional UMI-based ligation approaches such as those employed by anchored multiplex amplification [11] and single primer extension [12].

**Figure 1.**
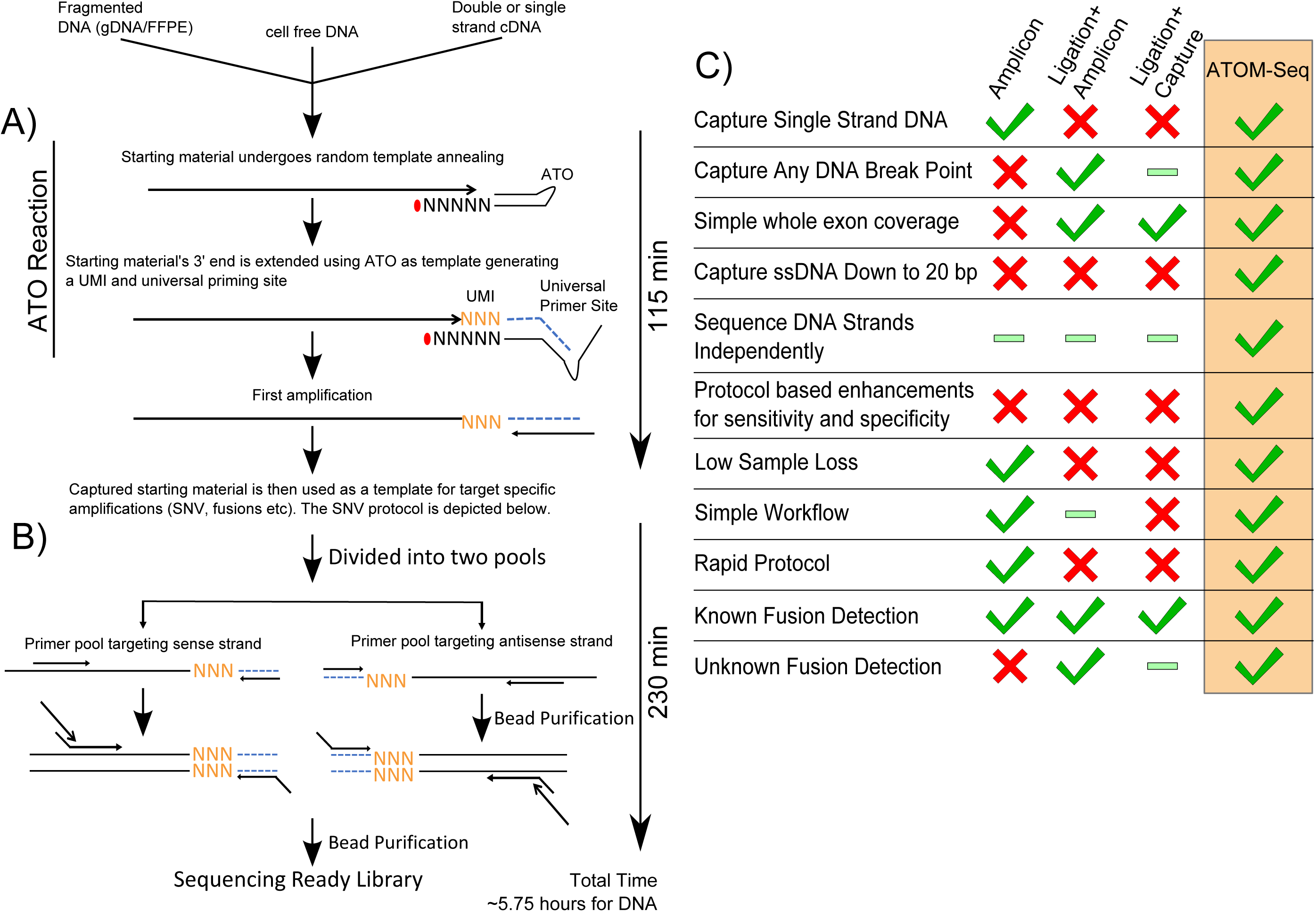
Overview and advantages of ATOM-Seq targeted enrichment protocols. ATOM-Seq based protocols can take any suitable stating material including cell free DNA, fragmented FFPE DNA, or cDNA. **A)** The first step is an “ATO-Reaction” which captures the starting material extending their 3’ ends using an ATO as a template generating a Unique Molecular Identifier and a universal primer site. **B)** Target enrichment is accomplished by dividing the first amplification into two separate pools for two rounds of nested PCR. Each pool targets either the sense or antisense DNA strands allowing both strands to be independently enriched. **C)** ATOM-Seq has many advantages relative to current approaches and is the only technology capable of combining the advantages of both PCR- and ligation-based approaches. The ATO Reaction has the advantage of capturing any ss/ds-DNA 3’ end (down to 20bp) allowing for unknown fusion detection. The first amplification imparts a unique advantage to both sensitivity and specificity due to it being a linear amplification. The remainder of the protocol has minimal hands on time and bead purification steps resulting in a low loss simple rapid workflow. Green ticks indicate aspects the technologies can do. Green bars indicate aspects of the technologies which are inefficient. Red crosses indicate aspects the technologies can not do.

Following the capture of the starting material we designed an enrichment protocol which maximises both sensitivity and specificity. The captured material undergoes an initial amplification step, which uses a universal primer complementary to the universal priming site at the 3’ of each starting DNA molecule (Figure 1A). Repeated rounds of this linear amplification create multiple copies of each original starting molecule. After this point, separate workflows were designed depending on the purpose of the library preparation. The first workflow is for targeted enrichment of frequently mutated regions from DNA (Figure 1B); the second is for known and unknown fusion detection from RNA input (Supplementary Figure 1), the third is whole sample (a.k.a. whole genome) library preparation (Supplementary Figure 2). Together, the technological basis of an ATO-Reaction combined with these downstream protocols imparts a series of unique advantages. Importantly ATOM-Seq includes all advantages ascribed to either PCR- or ligation-based approaches, whilst avoiding their respective limitations, overviewed in Figure 1C.

For targeted PCR enrichment-based detection of mutations, the first amplification product is split into two halves, each acting as a template for separate pools of target specific primers. This design allows for two primer pools to independently amplify either the sense or antisense DNA strands of the original starting material, in combination with a universal primer, while eliminating the risk of opposing primers interfering with one another. Strand-specific targeting is possible as a target specific primer can only exponentially amplify a DNA molecule if it forms a correctly orientated pair with the universal primer. Following this first round of PCR is a second round of nested PCR enrichment, which also incorporates dual sample indices (Figure 1B).

The variation of this protocol for detecting the presence of known and unknown fusions from an RNA sample is very similar to the above, comprising of two rounds target specific enrichment using nested primers. In this instance however, cDNA is generated prior to the ATO-Reaction (Supplementary Figure 1A,B), and the first amplification product is not split between pools of target specific primers (Supplementary Figure 1C). The whole genome version of this protocol uses the first amplification product as input for a second ATO-Reaction (Supplementary Figure 2B), which incorporates a second universal priming site at the opposite end of the captured molecule, allowing global PCR amplification of captured molecules (Supplementary Figure 2C).

### Performance of SNV Detection Protocol with Mutation Reference Standards

In order to thoroughly evaluate the performance of the ATOM-Seq technology, a series of commercially available reference standards were used in combination with a large panel of approximately 1200 target enrichment primers designed to amplify approximately 575 regions across 100 genes frequently mutated in cancer. In order to assess sensitivity, standards with different expected allele frequencies (AF) and input quantities were used. A 5% AF standard was used with both 0.5 ng and 1.0 ng of material, a 1.3% AF standard was used with both 1.0 ng and 5.0 ng, and a 0.13% AF standard was used with 20 ng. An overview of the sequencing data and sensitivity is shown in Table 1 and Figure 2.

**Table 1.** Sequencing quality and sensitivity of ATOM-Seq SNV protocol on reference samples. Genome mapping rates were >99% for all samples.

|  | AF | 5.0% |  | 1.3% |  | 0.13% |
| --- | --- | --- | --- | --- | --- | --- |
|  | Input (ng) | 0.5 | 1.0 | 1.0 | 5.0 | 20.0 |
| Average Depth Per Primer | Sense DNA | 395 | 807 | 870 | 2193 | 9361 |
|  | Antisense DNA | 426 | 973 | 1147 | 2448 | 9759 |
|  | Combined | 820 | 1780 | 2017 | 4640 | 19119 |
| Average Primer Uniformity | Sense DNA | 95.7% | 96.1% | 96.4% | 96.8% | 96.5% |
|  | Antisense DNA | 94.0% | 94.3% | 94.7% | 94.3% | 95.1% |
|  | Combined | 94.9% | 95.2% | 95.6% | 95.5% | 95.8% |
| Average Primer On-Target Rate | Sense DNA | 84.5% | 85.6% | 84.4% | 83.9% | 84.4% |
|  | Antisense DNA | 90.8% | 90.6% | 91.1% | 90.5% | 89.6% |
|  | Combined | 87.7% | 88.1% | 87.7% | 87.2% | 87.0% |
| Reference Mutations Detected Across 3 Replicates |  | 100% | 100% | 90.2% | 100% | 94.0% |

**Figure 2.**
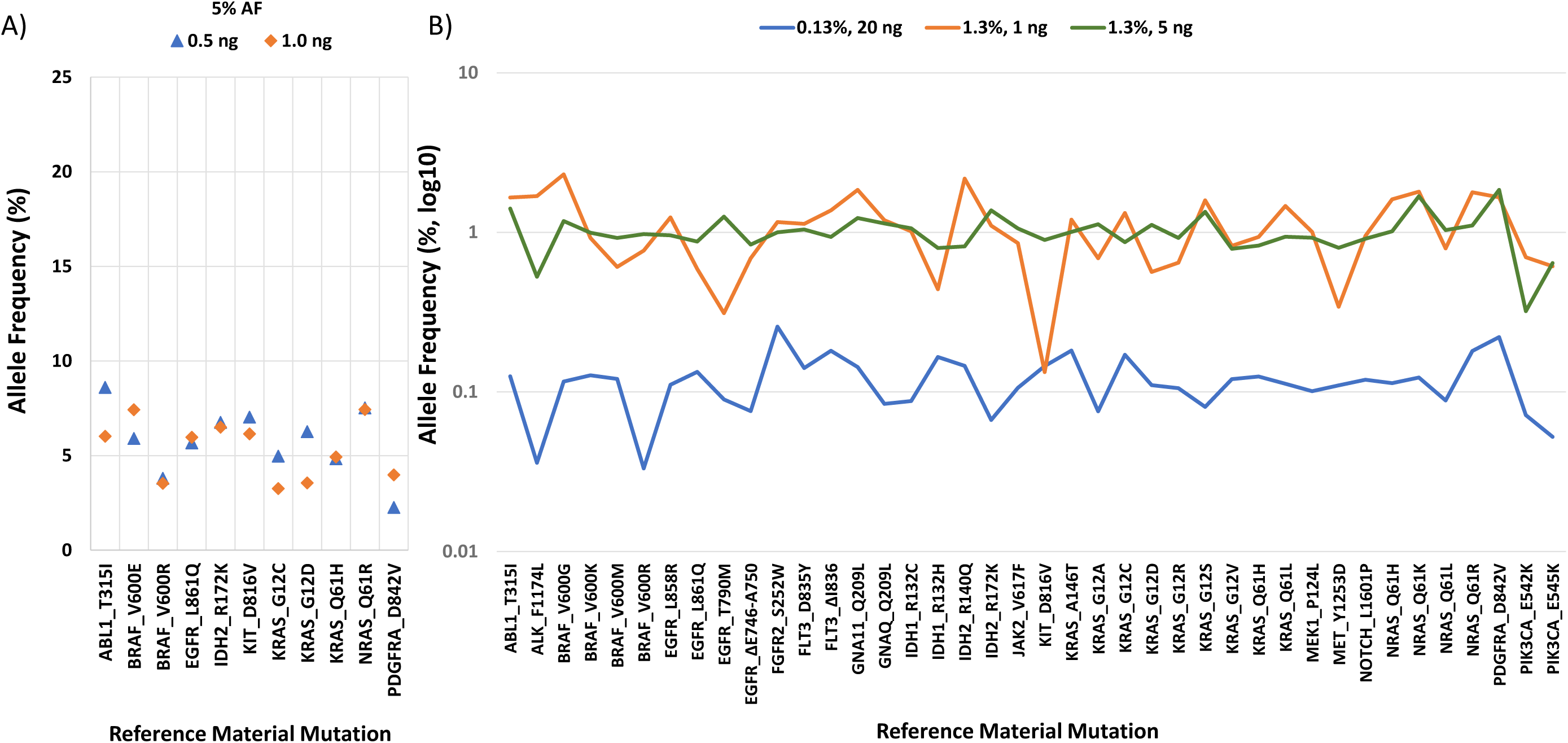
Technical validation of ATOM-Seq technology using reference standards. The plots show the average allele frequency (AF) across 3 biological replicates for each of the starting DNA quantities for **A) 5%** AF with 0.5 ng and 1 ng and **B) 1.3%** AF with 1 ng and 5 ng, and **0.13%** AF with 20 ng. The reference material contains variants with allele frequencies > 5% or >1.3%, which were detected but not shown.

With both 0.5 ng and 1.0 ng of reference material DNA containing 5% AF mutations a total of 100% of expected variants were detected in all three replicates. When using 1 ng of DNA with 1.3% AF, across all replicates 90% of mutations were found and between all replicates 100% of mutations were found. With 5 ng of DNA at 1.3% AF, 100% of the variants were detected in all 3 replicates. To test the absolute sensitivity of ATOM-Seq, the AF was reduced further to 0.13% and the input DNA quantity was increased to only 20 ng, proportionally 2.5x fewer copies of mutated material than used for 5 ng at 1.3% AF. Across all replicates 94% of mutations were found and between all replicates 100% of mutations were found (Figure 2).

The sensitivity and specificity of the ATOM-Seq protocol was examined with a semi-blind approach. First, we approximated the ‘ground truth’ by combining the three reference material replicates generated for 5 ng of starting material with 1.3% AF. With these combined samples we identified all variants which were identified with a count ≥3, this generated a list with between 149-153 variants across a total of 575 target regions per sample. Of these, a total of 149 were identified as ‘true positives’, those variants present in 2 or 3 of the 3 samples (Supplementary Figure 3). As summarised in Table 2, we found that true positive specificity was between 96.1-99.3% (average 97.2%) and true positive sensitivity was between 98.0-99.3% (average 98.7%).

**Table 2.** Sensitivity and specificity of ATOM-Seq SNV protocol on reference samples.

| Replicate | Variant Depth | Total Variants | True Positives Detected (n=149) | True Positive Specificity | True Positive Sensitivity |
| --- | --- | --- | --- | --- | --- |
| 1 | $\geq 3$ | 152 | 146 | 96.1% | 98.0% |
| 2 | $\geq 3$ | 153 | 147 | 96.1% | 98.7% |
| 3 | $\geq 3$ | 149 | 148 | 99.3% | 99.3% |
| Average |  | 151 | 147 | 97.2% | 98.7% |

### Performance of ATOM-Seq with FFPE and Liquid Biopsy DNA Samples

In order to clinically validate the ATOM-Seq technology, three small pilot studies were completed. These studies used cancer samples which had previously been assessed for mutations by one or more alternative approaches. Studies were performed blind with the expected results only shared upon completion of the sequencing analysis. An overview of all sequencing and identified variants is available in Supplementary Tables 1-4.

The first of these pilot studies used a colon and lung cancer panel of primers, targeting mutations common in these cancers. A total of 14 of 20 FFPE samples were successfully processed and sequencing. These samples had been previously assessed for the presence of *KRAS, BRAF, NRAS* or *PIKC3A* mutations using pyrosequencing, or for selected *TP53* mutations with targeted PCR and sequencing. Upon comparison of the data generated by ATOM-Seq to the expected data, all the previously identified mutations were identified. Further, due to the higher sensitivity of ATOM-Seq, two additional *PIK3CA* mutations were identified which were below the sensitivity threshold of pyrosequencing (Figure 3A).

**Figure 3.**
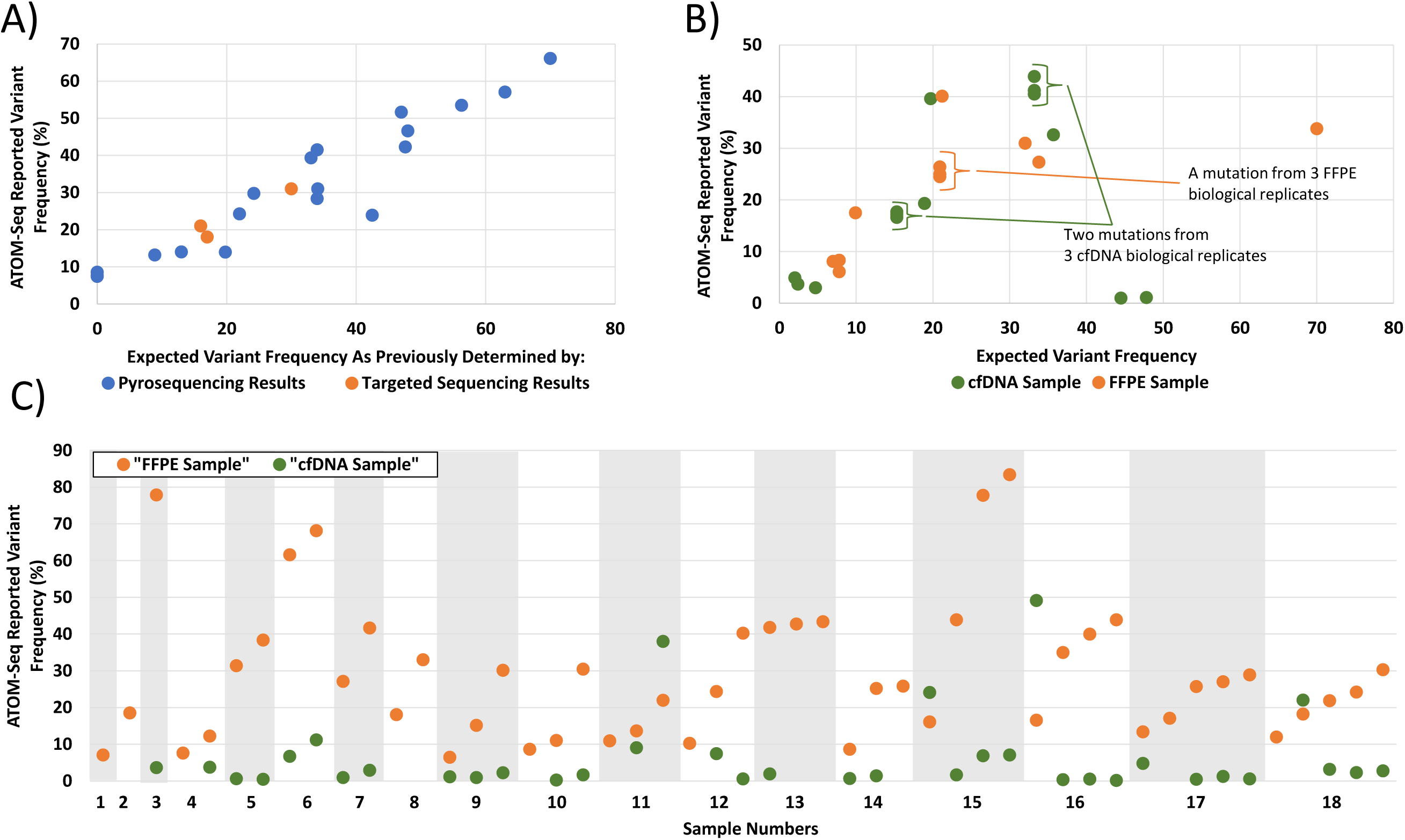
Technical validation of ATOM-Seq on clinical samples. **A)** Colon cancer samples. Correlation of the allele frequencies reported by ATOM-Seq relative to those determined either by pyrosequencing (blue spots) or targeted sequencing (orange spots). ATOM-Seq showed superior sensitivity by detecting variants in two samples which were not found when analysed by pyrosequencing as these were below the technology’s minimum sensitivity. **B)** Lung cancer samples. Correlation of the allele frequencies reported by ATOM-Seq relative to those determined by an alternative NGS approach. **C**) Paired lung cancer samples. Details of the allele frequencies for mutations found in FFPE samples and the concordant mutations from a matched cfDNA sample. A total of 84% (15/18) of patients had concordant mutations detected.

The second pilot study used the same panel of primers to identify mutations in unpaired lung cancer FFPE and cfDNA patient DNA samples. All samples were successfully processed, 7 cfDNA samples and 6 FFPE samples. From these, one sample of each material type was also processed in triplicate, to assess reproducibility. These samples had been previously sequenced using an alternative NGS technology, and all of the previously identified mutations were identified by ATOM-Seq. In the FFPE samples a total of 9 expected mutations were identified (AF 6.1-40.1%), with 10 expected mutations in the cfDNA samples (AF 1.0-41.2%). For those samples processed in triplicate, all mutations were observed at consistent allele frequencies (Figure 3B).

The third pilot study used a large 100 gene pan cancer panel. This was used for the identification of mutations from a series of 20 paired lung cancer FFPE and cfDNA samples, all samples were successfully processed and sequenced. No mutations were identified in two FFPE samples. For the remaining 18 paired samples we assessed the concordance between the FFPE and cfDNA samples. A total of 49 mutations were identified in the FFPE samples and of these 37 were identified in a paired cfDNA sample (Figure 3C). The allele frequency of the concordant mutations ranged between 0.13-49.1% for cfDNA and 6.38-83.3% for FFPE samples (Figure 3C). All 20 FFPE samples were previously sequenced using an alternative NGS technology. In genomic regions covered by both the ATOM-Seq panel and the previously used gene panels, all but 2 FFPE mutations were confirmed. A total of three of the cfDNA samples were also previously sequenced with only two concordant mutations detected, compared to eight concordant mutations detected using ATOM-Seq (samples 4, 9, and 18).

### Performance of ATOM-Seq with FFPE RNA and Liquid Biopsy Total Nucleic Acid Samples

Next, we evaluated the performance of an ATOM-Seq based protocol designed for detection of known and unknown fusions. To do this we used a commercial reference standard containing three validated fusions which we tested using a custom 170 primer panel designed to enrich target regions of cDNA known to frequently undergo fusion events in cancer. A total of 100 ng of FFPE RNA was used as starting material and the resultant library was sequenced to a depth of 3.1 million 150 bp PE reads. Upon analysis of the sequencing data the three expected fusion events were clearly identified (Figure 4A).

**Figure 4.**
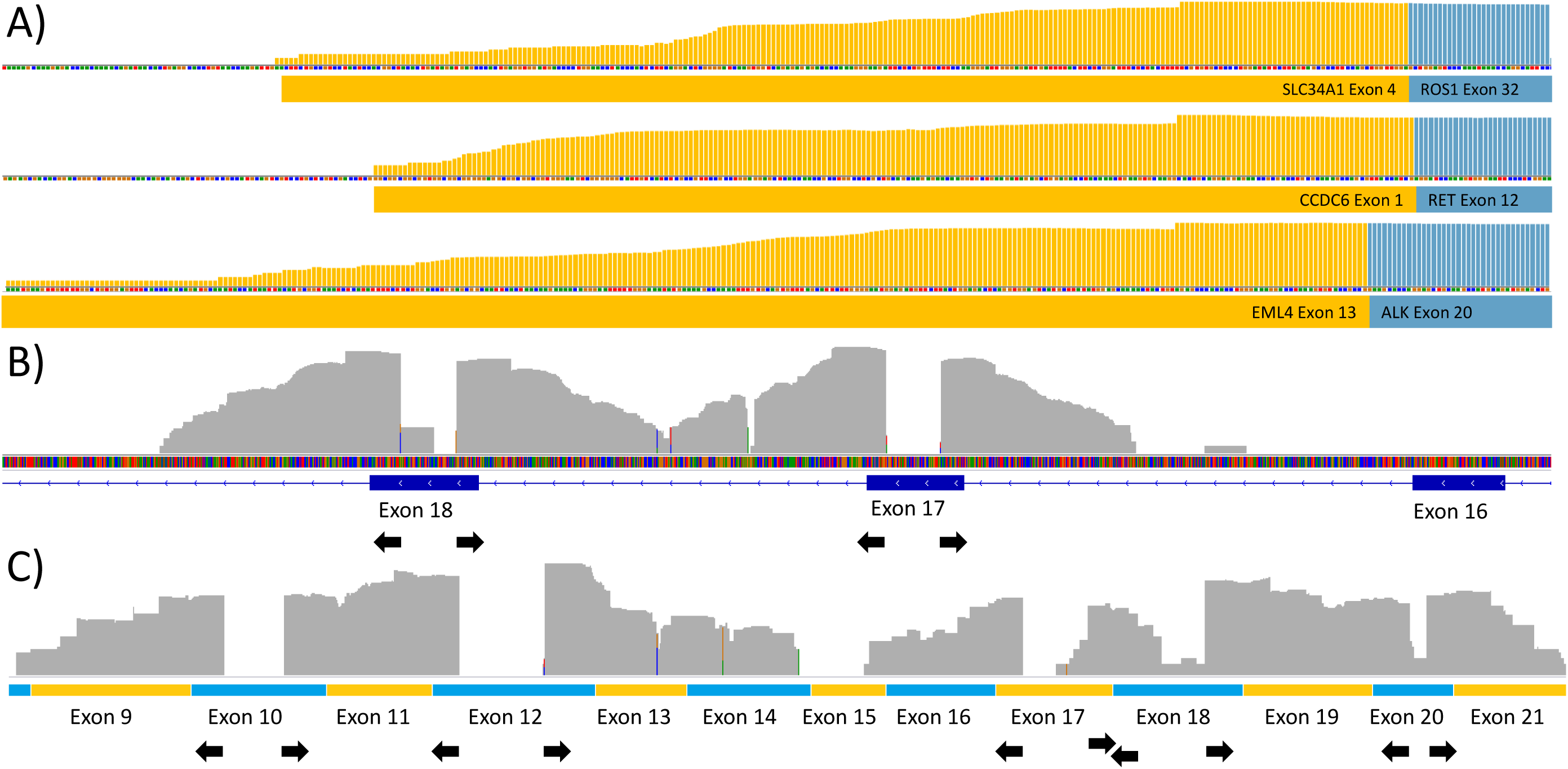
Technical validation of ATOM-Seq RNA based fusion protocol. **A)** An FFPE fusion reference material containing 3 certified fusions was used to demonstrate fusion detection. Reads were mapped to contigs representative of the fusions, demonstrating reads spanning the EML4-ALK, CCDC6-RET, and SLC34A2-ROS1 fusions. **B**,**C)** A total nucleic acid sample was used as stating material to the fusion protocol. **B)** Shows DNA derived reads mapped to hg38 spanning exon-intron boundaries. **C)** Shows reads which map to JAK1 (NM_001320923.1). The exons and orientation of the primers are shown, this demonstrates successful capture of cell free RNA molecules as shown by the exon junction spanning reads. All read depths are shown in a log10 scale.

To expand further we tested this protocol with a liquid biopsy total nucleic acid (TNA) extraction. With 13.65 ng of TNA we were able to successfully capture both DNA and RNA molecules from within a single TNA extraction (Figure 4B,C). As RNA constitutes a small portion of the nucleic acids present in an extraction, the majority of the reads can be assigned to DNA; these are identifiable as spanning intron-exon boundaries. To identify RNA derived reads, cDNA contigs were used for mapping and doing so allowed for clear identification of intron spanning reads indicating successful capture of cfRNA.

### Performance of ATOM-Seq Whole Genome Protocols

The ATOM-Seq technology was designed to be highly versatile, as such we tested its performance on a number of challenging and high-quality materials. Three different starting materials were used: cfDNA; fragmented genomic DNA; and single strand DNA PCR oligos as an analogue for ultra-short fragmented DNA, with a size distribution between 20-30bp. With cfDNA and fragmented genomic DNA the protocol was able to successfully capture both starting materials, in each case generating final libraries with profiles representative of their pre-capture profiles (Figure 5A-B). With the ultra-short DNA, upon mapping to the human genome, short 20-30bp stacks of reads were visible at all the expect primer sites with sizes down to 20bp represented in the data (Figure 5C).

**Figure 5.**
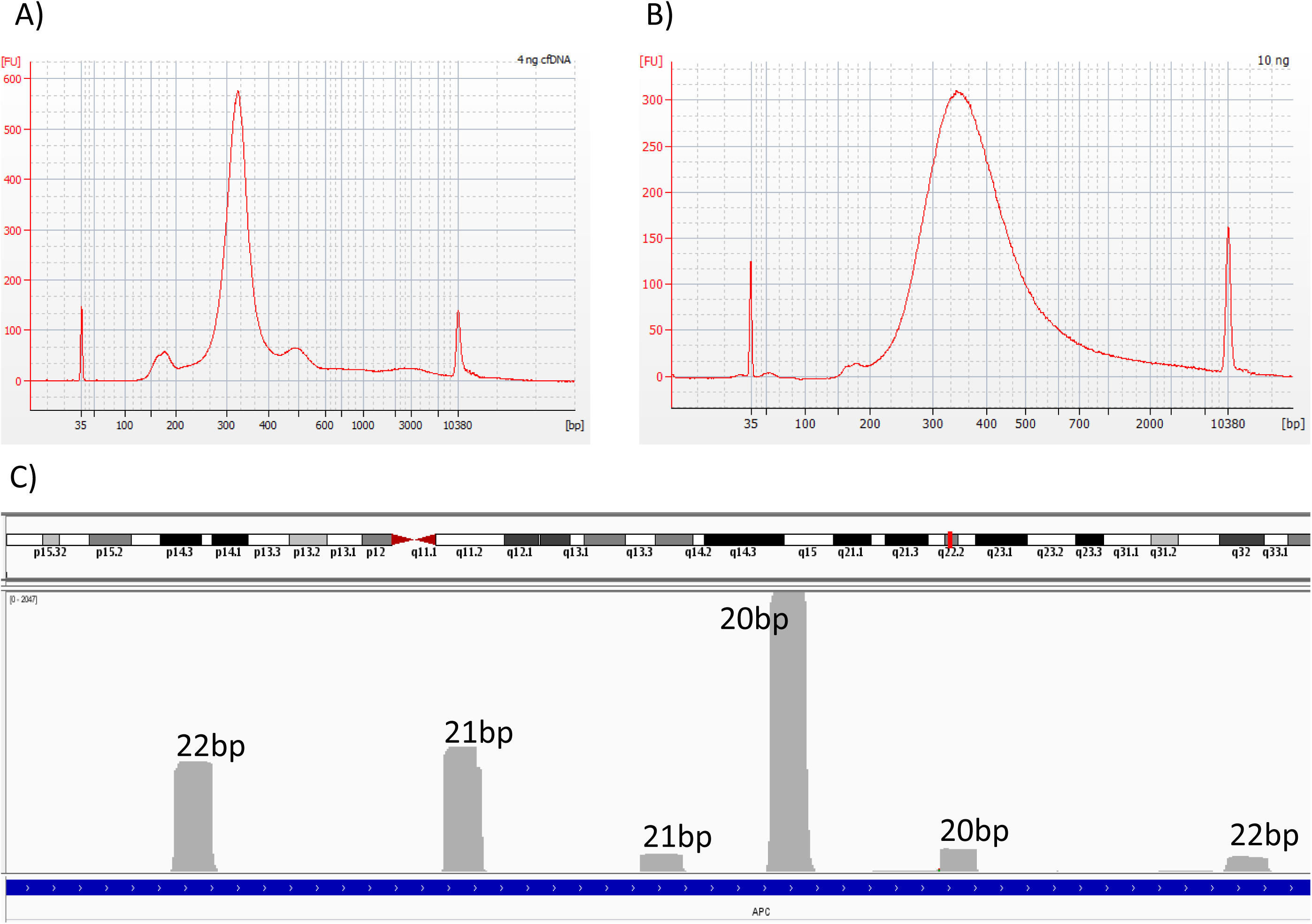
Results of whole sample library preparations using a series of different input materials. **A)** shows the final library profile generated when using 4 ng of cell free DNA, and **B)** shows the final library profile generated using 10 ng of fragmented gDNA. Both of these profiles mimic the respective size distribution profiles of their original starting materials. **C)** Shows mapped sequencing reads when very short DNA oligos were used as starting material. The width of the stacks are indicated individually. These correspond exactly to the length and expected genome mapping position of the oliogs used as starting material, demonstrating that capture of molecules as short as 20bp is possible using this method.

## Discussion

In recent years, many library preparation approaches have been adapted to testing of clinical material isolated from liquid biopsies, and they have improved the specificity of the data produced by reducing the background error rate of these approaches [8-13]. However, current technologies have inherent limitations which cannot be overcome due to the nature of PCR and ligation-based approaches. Because of this, new library preparation approaches are needed to ensure that clinical testing is able to assess the full spectrum of nucleic acids present in a liquid biopsy sample, including single stranded, short, and damaged molecules.

In this paper we have described a new technology which is uniquely suited to capturing nucleic acids extracted from a liquid biopsy; **<u>A</u>**daptor **<u>T</u>**emplate **<u>O</u>**ligo **<u>M</u>**ediated **<u>Seq</u>**uencing - ATOM-Seq. This technology is a highly robust, flexible and rapid approach for the generation of NGS libraries using DNA, RNA, or DNA+RNA mixtures as starting material. We have demonstrated its high sensitivity and specificity for detection of clinically relevant mutations (including SNV, deletions/insertions) from reference materials and its utility with clinical material, showing excellent performance with both FFPE DNA and cfDNA samples. As the technology is based on oligo annealing and polymerase driven extensions, which form the basis of all PCR reactions, it is naturally a highly efficient and effective process. The nature of the ATOM-Seq technology and its uniquely designed protocol imparts numerous significant advantages, as outlined in Figure 1C. These are all founded on its ability to capture the 3’ ends of any available nucleic acid, including double- and single-strand DNA, enzymatically or sonicated FFPE derived DNA, cfDNA, and single- or double-strand cDNA derived from total RNA (high quality or FFPE), cfRNA or cfDNA+cfRNA from a total cell free nucleic acid extraction. As any 3’ end can be captured, the technology will also be resistant to highly damaged and fragmented DNA. Combined, these factors mean ATOM-Seq is highly efficient at capturing a much greater breadth of starting material which cannot be captured by using only a traditional PCR-only, ligation + PCR or ligation + hybrid capture based approach. This makes ATOM-Seq the ideal technology for use with clinical samples, especially liquid biopsy samples.

ATOM-Seq’s unique advantages are also present in the workflow after the initial DNA capture. The most substantial of these is the first amplification, which is a linear amplification generating multiple copies of each original starting molecule. This is a central part of the protocol for two reasons: firstly it allows for greater protocol sensitivity; it enables the amplification product to be split between multiple downstream PCR reactions without sacrificing sensitivity, as all original molecules are represented in every reaction. Without this prior amplification, splitting a sample between multiple downstream amplifications would incur an immediate 50% drop in maximum sensitivity for each target primer, such as with Qiagen’s GeneRead QIAact AIT DNA UMI Panel. Safely dividing the sample enables the ability of utilising multiple primer pools for independent targeting of both the sense and antisense DNA strands, as well as allowing for comprehensive and simple dual strand whole exon coverage. This is important when detecting mutations in tumour suppressors such as *TP53*, something which PCR-based approaches inevitably struggle with due to limitations with tiling primers [8]. This approach also avoids the risk of opposing primers forming non-specific PCR products or otherwise interfering with one another if they were to be within a single reaction, improving the efficiency of the PCR reactions. This is a particular risk in heavily tiled regions such as would be necessary for the last, long exon of the *APC* gene. Smaller primer pools also benefit from greater PCR efficiency, meaning that PCR cycle number and protocol time can be kept lower.

The second advantage of incorporating a linear amplification is that it aids in the reduction of the error rate in the final library by improving the reliability and power of UMIs. In approaches where UMI incorporation is followed immediately by PCR amplification, any polymerase error that were to occur in the first round of target enrichment PCR would propagate exponentially and form a substantial fraction of the final PCR products [20]. The rounds of linear amplification in the ATOM-Seq protocols mean any randomly generated errors would be present in a far smaller fraction of final PCR products, reducing the chance that these are still present following error correction. Having the genetic information for both sense and antisense strands also allows for the comparison of mutations between strands, which increases confidence in mutation calling.

Current technologies have not, or cannot, incorporate this linear amplification and sample division method. PCR-based protocols incorporate UMIs in the first few rounds of PCR [8]. As such any attempt to incorporate a linear amplification would result in generation of errors which could not be differentiated from real mutations, defeating the purpose of UMI incorporation entirely. Ligation + PCR-based technologies such as Anchored Multiplex PCR or Single Primer Extension [11-13], apart from the aforementioned protocol, do not divide their samples for independent strand targeting. As they have their universal primer sites on the 5’ ends of DNA, not the 3’ as in ATOM-Seq, this means that linear amplification would have to be performed using their pools of targeted specific primers. Primer pools containing opposing pairs of primers would, 1) introduce bias due to variation in primer efficiencies and 2) inevitably pair together to produce PCR products, at the detriment of the protocol. Ligation + capture-based protocols could support linear amplification as they create 3’ universal primer sites, but doing this would add complexity to protocols which are already long.

The aforementioned advantages of ATOM-Seq bear through to the performance of the technology with the different types of reference and clinical materials. The presented data demonstrate the reliable detection of ≥5.0% AF mutations with as little as 0.5 ng of reference material and with between 10-40 ng of FFPE material. Not only does ATOM-Seq perform very well with this material, but when the identified mutations from clinical samples were compared with the allele frequencies determined by other technologies (pyrosequencing, ligation+PCR NGS, or PCR NGS) there was a very strong correlation between these data. This demonstrates that despite ATOM-Seq being a relatively young and unique technology it performs very well when compared with other much more mature technologies. Though ATOM-Seq was developed with liquid biopsy testing in mind, it is important to note that it also performs very well with FFPE material; formalin fixation still forms a central step in oncology patient sample processing worldwide.

We also demonstrate achieving reliable ≥0.1% AF mutation detection when testing reference material designed to be representative of cfDNA. While using liquid biopsy extracted material we were able to identify concordant mutations down to 0.12% in cfDNA, and in this pilot study generated a high level of concordance between paired FFPE and cfDNA samples, 15 of 18 paired samples. In a limited sample set (n=3) ATOM-Seq also outperformed a ligation-based NGS technology by detecting more cfDNA mutations: 8 versus 2. This can in part be attributed to ATOM-Seq’s ability to capture a greater proportion of cfDNA which is enriched for tumour derived material (short and single stranded DNA fragments) given the equivalent depths between technologies at these sites (Supplementary Table 4). ATOM-Seq was also able to identify all expected types of mutations from paired FFPE and cfDNA clinical material, including SNPs, deletions as long as 28bp (found in *BRCA2*) and insertions up to 9bp (found in *ERBB2;* Supplementary Table 4).

Not only do ATOM-Seq-based approaches contain many inherent advantages, but targeted enrichment from cfDNA to purified final library can be accomplished in only 5.75 hours, making it an ideal protocol to complete in less than one day (Figure 1). When using RNA, proceeding from RNA/cfRNA/TNA to a bead purified final library can be completed in just over 7 hours, again making it a less than one day protocol (Supplementary Figure 1). Whole genome library preparation can be completed in just over five hours (Supplementary Figure 2). ATOM-Seq protocol are more time efficient when compared to ligation based protocols, which can take from eight hours for ligation+PCR protocols to up to three days for ligation+capture protocols [8-13].

ATOM-Seq’s excellent performance with both FFPE material and especially with cfDNA demonstrates that it would be an ideal technology for incorporation into both research and clinical oncology sample testing environments. The combination of the underlying approach, streamlined flexible protocols, and technological advantages (including the unique demonstrated ability to capture ssDNA down to 20bp) establishes ATOM-Seq as perfectly and uniquely suited for adoption into testing for cancer associated mutations using challenging liquid biopsy material.

## Methods

### Preparation of Starting Materials

Reference material was purchased from Horizon Discovery, Tru-Q 4 (HD731) containing 5% allele frequency mutations, Tru-Q 7 (HD734) containing 1.3% allele frequency mutations, and a wild type sample Tru-Q 0 (HD752). In order to create a 0.13% allele frequency sample, the 1.3% material was diluted 1:10 in the WT material. To create a fragmented DNA analogue with a size distribution similar to that of cfDNA, the 5%, 1.3%, and 0.13% samples were independently fragmented using the KAPA Frag kit (Roche; 07962517001) following the manufacturers recommended conditions followed by a 3x AMPure bead purification. FFPE samples were either enzymatically fragmented as above or sonicated using a Covaris ultrasonicator. The size distribution was measured using an Agilent Bioanalyzer High Sensitivity kit and concentrations were determined using a Qubit dsDNA HS Assay Kit (Thermo Fisher Scientific; Q32854).

Fusion reference material was purchased from Horizon Discovery, an FFPE sample containing 3 validated fusions (HD784). RNA was extracted using a ReliaPrep™ FFPE Total RNA Miniprep System kit (Promega; Z1001) following the manufacturers recommended protocol. Frozen serum was purchased Cambridge Bioscience Ltd (PLSSKF8BC200-XSXX) and total nucleic acids were extracted using MagMAX™ Cell-Free Total Nucleic Acid Isolation Kit following the manufactures recommended conditions (Applied Biosystems; A36716)

### Primer Panel Design

A Pan Cancer hotspot panel containing the most frequently mutated sites across a broad range of cancers was designed by examining COSMIC database version 85. The Pan Caner panel was used while testing technical sensitivities on reference materials. The primer sequences were designed taking into account coverage of target sites, balancing primer Tm, and predicted specificity within the human genome (HG38). The final full pan cancer panel includes 100 genes which cover approximately 4,000 annotated COSMIC mutations. There are 577 primer sites targeting the sense DNA strand and 570 primer sites targeting the antisense DNA strand. All enrichment amplifications are designed to have two rounds of PCR, and as such an ‘outer’ and ‘inner’ primer were designed for all target sites, where the inner primers contain a universal tail to allow for preparation of dual index libraries. All primers were synthesised by Eurofins Genomics. A smaller 23 gene panel was created as a subset of the pan cancer panel for specifically testing colon cancer and lung cancer cfDNA and FFPE clinical samples. The smaller panel has 131 primers targeting the antisense DNA strand and 132 targeting the antisense DNA strand. Fusion Panel Design followed an equivalent process as the pan cancer panel except for targeting gene fusions frequently detected in cancer. Placement of enrichment primers in a conserved fusion partner allows for detection of unknown fusion partners. The final panel targeted 169 conserved exons found as fusion partners across 22 genes. Details of all target genes are in Supplementary Table 5.

### ATO-Reaction – 3’ Extension Reaction

For reference material testing, different quantities of fragmented DNA were used in triplicate for each allele frequency being tested. For 5% allele frequency either 0.5 ng or 1.0 ng was used, for 1.3% either 1.0 ng or 5.0 ng was used, and for 0.13% 20 ng was used. When testing fusion detection with reference material, a total of 100 ng of RNA was used. For TNA a total of 13.65 ng was used as determined by DNA mass, measured using a Qubit 3.0. For clinical material various quantities were used. For the first pilot study 10 ng of Covaris sheared FFPE extracted DNA were used. For the second pilot study 40 ng of enzymatically fragmented FFPE and 11.8-20.96 ng of cfDNA were used. For the third pilot study 40 ng of enzymatically fragmented FFPE DNA and 25-64 ng of cfDNA were used.

All methods (depicted in Figure 1A, sup Figure 1, 2) were performed as described in the XCeloSeq Pan Cancer Kit (GeneFirst; SEQ002), XCeloSeq cfDNA Library Prep Kit (GeneFirst; SEQ001), or XCeloSeq Fusion Research Kit (GeneFirst; SEQ007) following the manufacturer’s recommended protocol.

Briefly, when XCeloSeq Fusion Research Kit and RNA was used, first strand cDNA was made by mixing 4 μl of ‘FS Mix’, RNA, and nuclease free water to a final volume of 20 μl and thermocycling as follows: 65°C for 5 min, 4°C for 2 min, 25°C for 2 min, and 55°C for 10 min. To this, 1.5 μl of ‘SS Enzyme’ was added and incubated at 22°C for 30 min. Second strand cDNA was purified with AMPure XP beads with a 2.0x ratio and eluted into 13 μl nuclease free water.

The following steps vary slightly depending on the samples and version of the protocol used. All samples followed the same general procedure. Briefly, the XCeloSeq Pan Cancer Kit and XCeloSeq cfDNA Library Preparation kit protocols are as follows. The cfDNA, fragmented FFPE DNA, freshly made cDNA, or DNA oligos were combined with 2 μl of ‘Adaptor Template Oligo (ATO)’ to a final volume of 15 μl for the Pan Cancer Kit, or, 1 μl of ATO to a final volume of 7.5 μl for the cfDNA Kit. This mixture was heated to 65°C for 2.5 min before being cooled to 4°C. To this 3 μl of ‘ATO Reagents’ and 2 μl of ‘ATO Enzyme’ are added for the Pan Cancer Kit or 1.5 μl and 1 μl respectively for the cfDNA Kit. To induce the extension of the starting material as a primer using the ATO as a template the mixture is placed in a pre-cooled 4°C thermocycler and cycled as follows, 26°C for 6 min, 30°C for 10 min, 65°C for 1 min, 10°C for 1 min, 26°C for 6 min, 30°C for 10 min followed by two cycles of 65°C for 1 min, 10°C for 1 min, 26°C for 6 min and 30°C for 5 min. For the Pan Cancer Kit 25 μl of ‘Amplification One Mix’ and 1 μl of ‘Primers’ are combined and cycled as follows 37°C for 10 min, 98°C for 30 sec followed by 10 cycles of 98°C for 5 sec, 60°C for 1 min, and 72°C for 1 min and with a final extension of 72°C for 2 min. For the cfDNA Kit 1 μl of ‘ATO Treatment’ is added and incubated for 37°C for 20 min and 25°C for 10 min. To this product 1.5 μl of ‘Amplification Primers’ and 12.5 μl of ‘Universal Enzyme Mix’ were combined and cycled as follows 98°C for 30 sec followed by 6 cycles of 98°C for 10 sec, 65°C for 75 sec, and with a final extension of 65°C for 2 min. The product of this is the ‘First Amplification Product’ which is used for downstream target enrichment.

### ATOM-Seq Targeted Hot Spot Enrichment

For the first PCR in the Pan Cancer Kit, 25 μl of ‘Master Mix’, 3 μl ‘Pool 1 – OUTER’ or ‘Pool 2 – OUTER’ primer mixes targeting the sense or the antisense DNA strands and 22 μl of the First Amplification Product were combined. Both of these mixtures were thermocycled as follows, 98°C for 30 sec followed by 14 cycles of 98°C for 5 sec, 65°C for 5 min, and 72°C for 30 sec followed by a final extension of 72°C for 2 min. These Second Amplification Products were purified with AMPure beads with a 1.8x ratio. For the second PCR, 25 μl ‘Master Mix’, 2 μl of ‘Pool 2 – INNER’ or ‘Pool 2 – INNER’ nested primer mixes targeting the sense or the antisense DNA strand, 1 μl of an i7 index primer and 1 μl of an i5 index primer and 21 μl of the appropriate bead purified Second Amplification Product were combined. Both of these mixtures were thermocycled as previously. Finally, both samples were purified with AMPure beads with a 1.2x ratio.

### ATOM-Seq Targeted Fusion Enrichment

For the XCeloSeq Fusion Research Kit the first PCR differs in that there is only a single pool of primers. As such, 2.5 μl of primers and an additional 1.5 μl of master mix were added directly to the entire 46 μl First Amplification Product, which was then cycled as per the Pan Cancer Kit. These were also bead purified and amplified again following the procedure for the Pan Cancer Kit. A schematic of the protocol is detailed in Supplementary Figure 1.

### ATOM-Seq Whole Genome Protocol

The ‘whole genome’ library preparation was performed using the XCeloSeq cfDNA Library Preparation kit (GeneFirst; SEQ001). The first ATO-Reaction and First Amplification were performed as detailed above and this was then followed by a bead purification step, as detailed in Supplementary Figure 2. Following this a second ATO-Reaction was preformed using the First Amplification product as a template. The steps are equivalent to the first ATO-Reaction except the cycling was as follows: 26°C for 6 min, 30°C for 10 min, 65°C for 1 min, 10°C for 1 min, 26°C for 6 min, 30°C for 10 min. Finally a global sample amplification using i5 and i7 index primer was used to produce a final sequencing library cycled as follows 98°C for 30 sec, 8 cycles of 98°C for 10 sec 60°C for 30 sec, 65°C for 75 sec followed by a final incubation of 65°C for 2 min.

### Sequencing and Data Analysis

Sequencing was generated using either a MiSeq or NextSeq. High depth sequencing was generated using a HiSeq at the Wellcome Centre for Human Genetics, Oxford. Reference material sequencing data was generated using a MiSeq. For the first pilot study each sample was sequenced to a depth between 0.7-5.1 million 150 bp PE reads on a MiSeq. For the second pilot study each FFPE and cfDNA sample was sequenced with between 4.6-5.6 million 150 bp PE reads on a single NextSeq run. For the third pilot study each FFPE sample was sequenced with between 3.3-79 million 150 bp PE reads and each cfDNA sample between 2.9-79 million 150 bp PE reads on either a MiSeq or HiSeq.

Sequencing data was processed using the following steps. Initially, adaptors were trimmed from reads, 20 bp UMIs were extracted from the beginning of read 2 into read headers, and inserts below 30 bp were discarded using fastp v0.20.1 [21] with default settings alongside custom python scripts. The trimmed reads were then mapped to the HG38 reference genome using BWA v0.7.15 [22] with default settings. Mapped reads were passed to gencore v0.15.0 [23] for consensus read generation and error correction using default settings (80% consensus to generate a consensus base) with a minimum family size of two. Consensus reads were parsed by a custom python script which removed any reads not mapping to a primer site and trimmed the primer sequence from those reads which did. Variants were then called by passing the filtered and trimmed consensus reads to VarDict v1.7 [24].and calling variants only within the targeted regions using the following settings -f 0.0001 -r 1 - M 10 -P 0 -Q 10 -U -u. Variant lists were then filtered further to remove variants with AF <5.0 and counts <3 from FFPE samples along with additional manually inspecting consensus reads in Integrative genomics Viewer [25].

To assess the specificity of the ATOM-Seq protocol we approximated the ‘ground truth’ from three reference material replicates. This was done by generating a combined list of variants across the three replicates generated using the 1% AF with 5 ng of starting material. A minimum variant count of 3 was used as a cut off, any variants below this threshold were treated as error and discarded. From this list ‘true positives’ were assessed to be those which were present in 2 or 3 of the 3 replicates (Supplementary figure 3).

All sequencing data generated using RNA or TNA was processed with the following pipeline. Raw reads were trimmed as before, mapped to the HG38 reference genome using the splice aware mapper STAR [26] or to cDNA contigs using BWA, and then reference fusions were identified from the mapping output. Custom fusion contigs were used to accurately depict fusion spanning reads (COSF408 (EML4-ALK); COSF1271 (CCDC6-RET); COSF1196 (SLC34A2-ROS1)). For cell free total nucleic acids, cfDNA derived reads were identified based on the reads spanning intron/exon boundaries, cfRNA derived reads were identified being exon junction spanning after mapping to JAK1 mRNA contigs (NM_001320923.1).

## Acknowledgements

We thank the Oxford Genomics Centre at the Wellcome Centre for Human Genetics (funded by Wellcome Trust grant reference 20314/Z/16/Z) for the generation and initial processing of the HiSeq sequencing data presented herein. We thank Jingchen Chen for his input with technology development, primer design, sample processing and data analysis.

The work in this paper was in part funded by Innovate UK Open Programmes Round 3 (project ID 1050), Innovate UK Shanghai-UK Industrial Challenge Programme (project ID 98238-574182), and the Shanghai International Science & Technology Cooperation Fund (17440732400).

## Author Contributions

G.F. conceived the study. G.F and T.L.D. participated in project design. T.L.D., S.C.D., J.Y, P.W.B. and S.S. participated in technology development and performed experimental work. T.L.D completed reference material data and clinical sample data analysis. S.R., H.W., H.S. and D.B. performed colorectal cancer sample processing and analysis. A.L.O., S.G.F.W., B.H.G. and H.Y.D. performed paired lung cancer sample processing and analysis. X.Y. and H.X. performed lung cancer sample processing and analysis. T.L.D., S.C.D. and G.F. wrote, reviewed and approved the manuscript. All authors approved the submitted version of the manuscript.

## Competing interests

G.F., T.L.D., S.C.D., J.Y, P.W.B. and S.S. are or were direct employees of GeneFirst. X.Y. and H.X. are or were employees of Guangzhou Biotron Technology Co., Ltd which has financial interest in GeneFirst. All other authors declare no conflicts of interest.

## Figure Legends

**Supplemental Figure 1:** Overview of an ATOM-Seq Fusion enrichment protocol.

**Supplemental Figure 2:** Overview of an ATOM-Seq ‘whole genome’ protocol.

**Supplemental Figure 3:** Number of variants detect in each of three biological replicates with a variant count greater than 3.

**Supplemental Table 1:** Details of all sequencing for pilot studies 1, 2 and 3.

**Supplemental Table 2:** A list of variants from clinical samples for pilot study 1.

**Supplemental Table 3:** A list of variants from clinical samples for pilot study 2.

**Supplemental Table 4:** A list of variants from clinical samples for pilot study 3 using paired lung cancer samples.

**Supplemental Table 5:** A list of target genes present in all primer panels.

