## Supplemental Figures and Tables for "Adaptor Template Oligo-Mediated Sequencing (ATOM-Seq): A versatile and ultra-sensitive UMI-based NGS library preparation technology, for use with cfDNA and cfRNA"

### Supplementary Figures

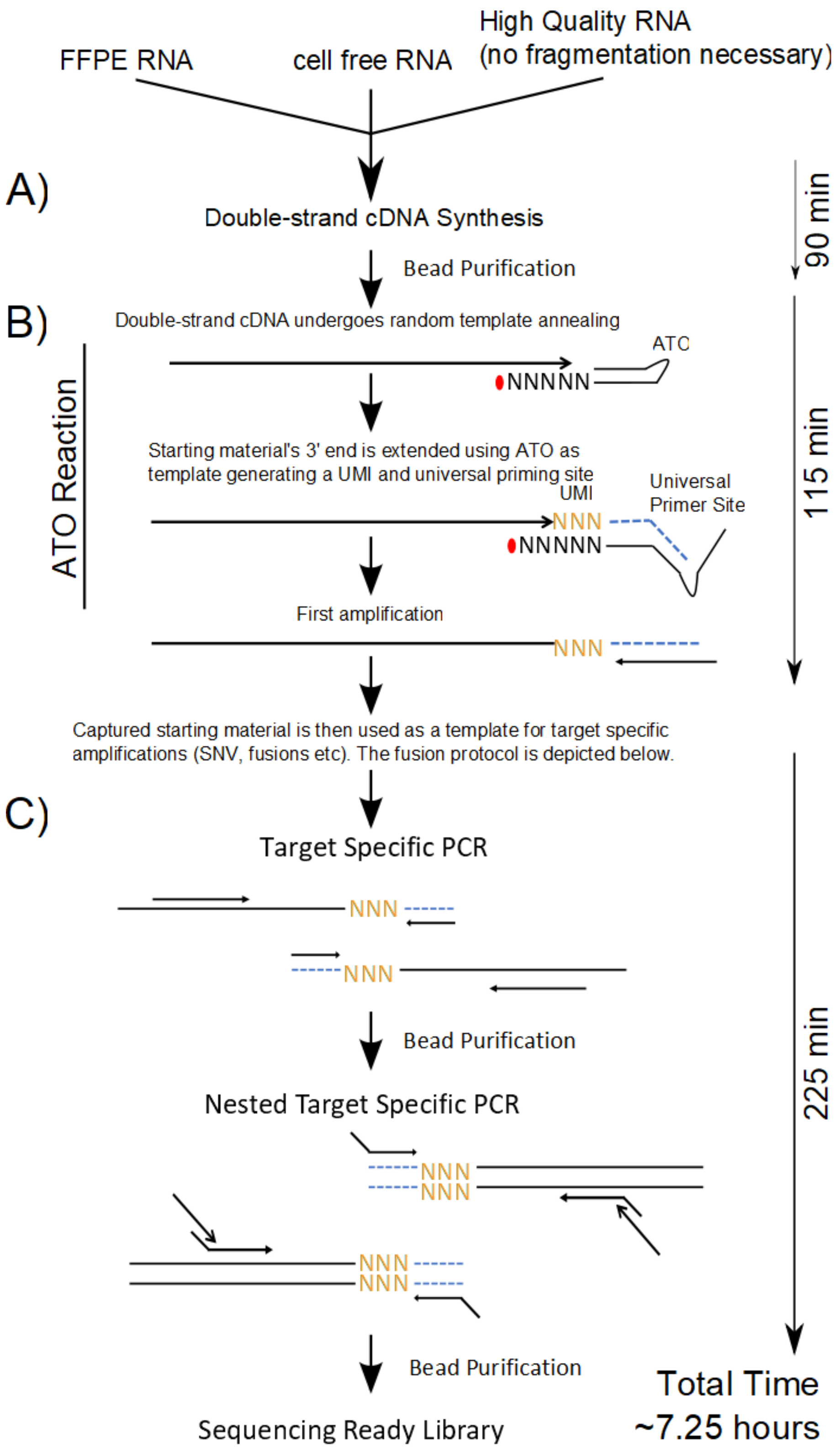

**Supplementary Figure 1.** Overview of an ATOM-Seq Fusion enrichment protocol. **A)** The first step is double strand cDNA synthesis followed by a bead purification. **B)** An “ATO-Reaction” then captures the ds-cDNA and 3’ ends are extended by a polymerase, following which is an initial round of amplification. **C)** The whole of the initial amplification is used in two rounds of nested PCR using target specific enrichment primers.

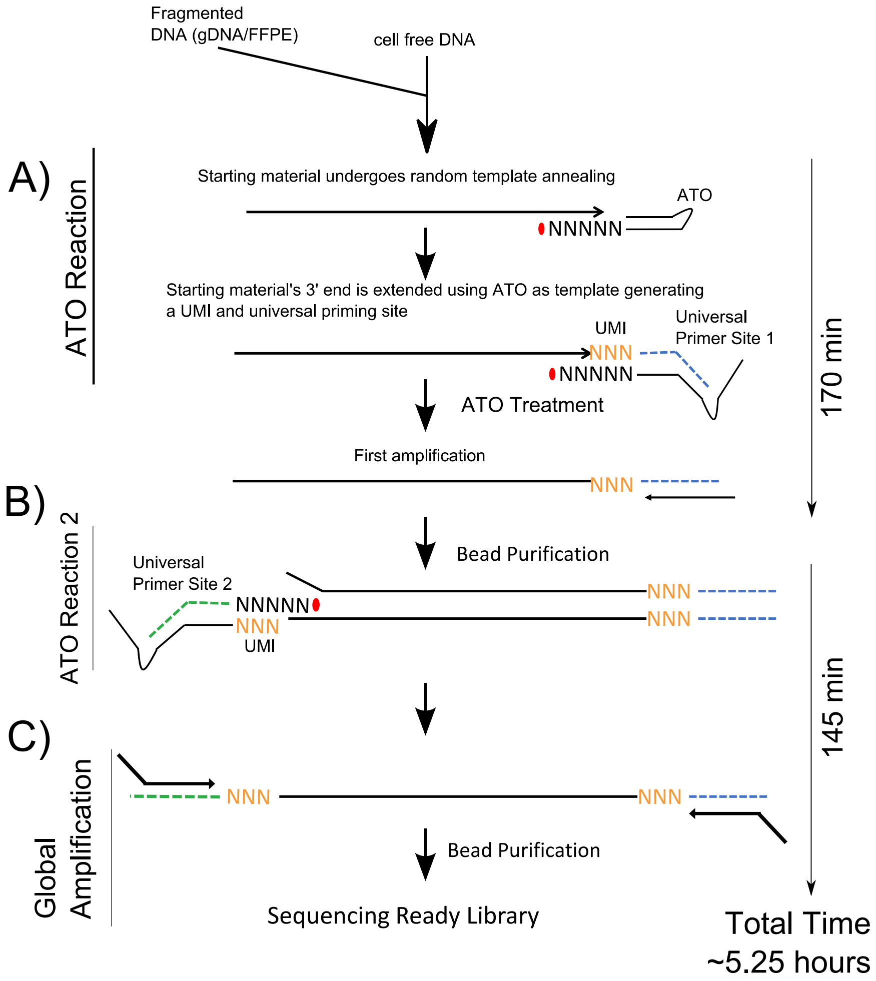

**Supplementary Figure 2.** Overview of an ATOM-Seq 'whole genome' protocol. ATOM-Seq based protocols can take any suitable stating material including cell free DNA, fragmented FFPE DNA, or cDNA. **A)** The first step is an "ATO-Reaction" which captures the starting material by annealing a synthetic Adaptor Template Oligo (ATO) to the 3' ends of all starting material. The 3' ends are then extended by a polymerase which uses the ATO as a template. This extension generates a Unique Molecular Identifier and a universal primer site on all 3' ends, following which is the first amplification. **B)** The first amplification product is used as an input for a second ATO-Reaction which creates a second, differing universal primer binding site. **C)** The two universal primer binding sites are used for whole library amplification.

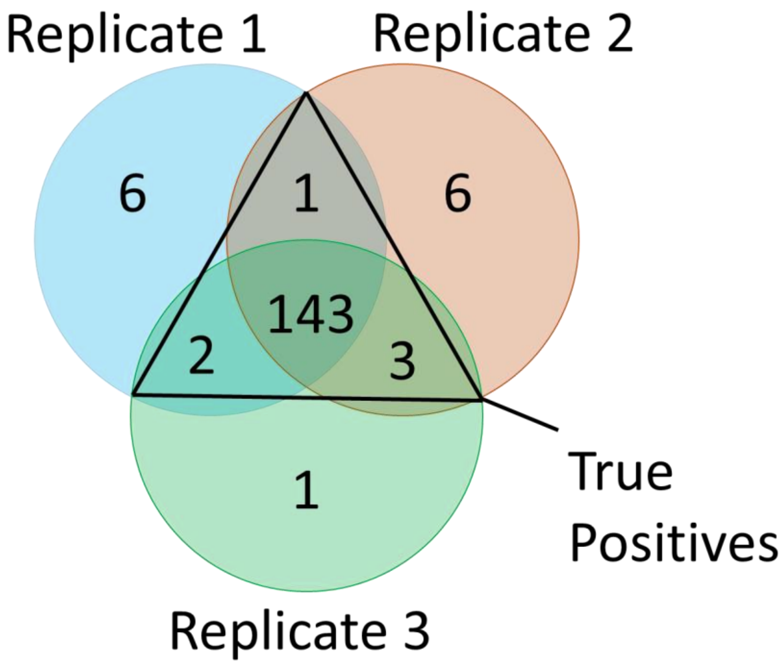

**Supplementary Figure 3.** Number of variants detect in each of three biological replicates with a variant count greater than 3. The variants are separated into those detected in only one sample, in 2 of 3 samples, or in all 3 samples. Those samples present in 2 or 3 of 3 samples were deemed 'true positives'

| Pilot Study | Sample ID | Cancer Type | Sample Type | Sequencing Instrument | PE Reads | Genome Mapping Rate (%) | Average Molecular Depth Across All Primers |
|---|---|---|---|---|---|---|---|
| 1 | 1 | Colon | FFPE | MiSeq | 785030 | 94.3 | 120 |
| 1 | 2 | Colon | FFPE | MiSeq | 1784784 | 99.5 | 350 |
| 1 | 3 | Colon | FFPE | MiSeq | 2559946 | 99.4 | 800 |
| 1 | 4 | Colon | FFPE | MiSeq | 869123 | 94.1 | 100 |
| 1 | 5 | Colon | FFPE | MiSeq | 714666 | 96.3 | 105 |
| 1 | 6 | Colon | FFPE | MiSeq | 849723 | 96.8 | 130 |
| 1 | 7 | Colon | FFPE | MiSeq | 803139 | 96.2 | 150 |
| 1 | 8 | Colon | FFPE | MiSeq | 966072 | 97.3 | 125 |
| 1 | 9 | Colon | FFPE | MiSeq | 925284 | 97.6 | 145 |
| 1 | 10 | Colon | FFPE | MiSeq | 957313 | 98.4 | 165 |
| 1 | 11 | Colon | FFPE | MiSeq | 1004834 | 97 | 130 |
| 1 | 12 | Colon | FFPE | MiSeq | 688384 | 89.3 | 120 |
| 1 | 13 | Colon | FFPE | MiSeq | 5074762 | 99.2 | 560 |
| 1 | 14 | Colon | FFPE | MiSeq | 948665 | 98.3 | 360 |
| 2 | 1 | Lung | cfDNA | NextSeq | 5588152 | 99.925 | 623 |
| 2 | 2 | Lung | cfDNA | NextSeq | 5823697 | 99.93 | 712 |
| 2 | 3 | Lung | cfDNA | NextSeq | 5667624 | 99.92 | 920 |
| 2 | 4 rep 1 | Lung | cfDNA | NextSeq | 6299592 | 99.95 | 1432 |
| 2 | 4 rep 2 | Lung | cfDNA | NextSeq | 5300468 | 99.935 | 1378 |
| 2 | 4 rep 3 | Lung | cfDNA | NextSeq | 5572667 | 99.95 | 1394 |
| 2 | 5 | Lung | cfDNA | NextSeq | 6076756 | 99.94 | 885 |
| 2 | 7 | Lung | cfDNA | NextSeq | 5815572 | 99.925 | 1272 |
| 2 | 8 | Lung | cfDNA | NextSeq | 5480830 | 99.95 | 1393 |
| 2 | 9 | Lung | FFPE | NextSeq | 5640270 | 99.92 | 2605 |
| 2 | 10 | Lung | FFPE | NextSeq | 5249362 | 99.9 | 2027 |
| 2 | 11 | Lung | FFPE | NextSeq | 4814414 | 99.865 | 1386 |
| 2 | 12 rep 1 | Lung | FFPE | NextSeq | 5908447 | 99.64 | 2320 |
| 2 | 12 rep 2 | Lung | FFPE | NextSeq | 6580624 | 99.92 | 2347 |
| 2 | 12 rep 3 | Lung | FFPE | NextSeq | 5254466 | 99.905 | 2200 |
| 2 | 13 | Lung | FFPE | NextSeq | 4954218 | 99.91 | 2355 |
| 2 | 14 | Lung | FFPE | NextSeq | 4644891 | 99.915 | 2584 |
| 3 | 1 | Lung | FFPE | HiSeq | 14602226 | 99.9 | 699 |
| 3 | 1 | Lung | cfDNA | HiSeq | 39846459 | 99.8 | 341 |
| 3 | 2 | Lung | FFPE | MiSeq | 7416038 | 99.47 | 502 |
| 3 | 2 | Lung | cfDNA | MiSeq | 3287358 | 99.49 | 318 |
| 3 | 3 | Lung | FFPE | MiSeq | 6725536 | 99.24 | 317 |
| 3 | 3 | Lung | cfDNA | MiSeq | 3368818 | 99.07 | 157 |
| 3 | 4 | Lung | FFPE | HiSeq + MiSeq | 58845179 | 99.92 | 1302 |

| | | | | | | | |
|---|---|---|---|---|---|---|---|
| 3 | 4 | Lung | cfDNA | HiSeq + MiSeq | 65825543 | 99.57 | 404 |
| 3 | 5 | Lung | FFPE | HiSeq | 13025985 | 99.82 | 736 |
| 3 | 5 | Lung | cfDNA | HiSeq | 40727884 | 99.91 | 402 |
| 3 | 6 | Lung | FFPE | MiSeq | 3388633 | 99.19 | 341 |
| 3 | 6 | Lung | cfDNA | MiSeq | 6993570 | 99.22 | 192 |
| 3 | 7 | Lung | FFPE | MiSeq | 6395028 | 99.25 | 710 |
| 3 | 7 | Lung | cfDNA | MiSeq | 3288506 | 99.22 | 322 |
| 3 | 8 | Lung | FFPE | HiSeq | 13399391 | 99.94 | 494 |
| 3 | 8 | Lung | cfDNA | HiSeq | 42230425 | 99.92 | 399 |
| 3 | 9 | Lung | FFPE | HiSeq + MiSeq | 68031679 | 99.72 | 1365 |
| 3 | 9 | Lung | cfDNA | HiSeq + MiSeq | 83936128 | 99.57 | 458 |
| 3 | 10 | Lung | FFPE | HiSeq | 14949016 | 99.89 | 686 |
| 3 | 10 | Lung | cfDNA | HiSeq | 40671065 | 99.91 | 419 |
| 3 | 11 | Lung | FFPE | HiSeq | 13136302 | 99.83 | 613 |
| 3 | 11 | Lung | cfDNA | HiSeq | 61735997 | 99.93 | 623 |
| 3 | 12 | Lung | FFPE | HiSeq | 16275543 | 99.94 | 569 |
| 3 | 12 | Lung | cfDNA | HiSeq | 30399924 | 99.6 | 280 |
| 3 | 13 | Lung | FFPE | HiSeq | 13354101 | 99.3 | 507 |
| 3 | 13 | Lung | cfDNA | HiSeq | 36594461 | 99.79 | 280 |
| 3 | 14 | Lung | FFPE | MiSeq | 8502361 | 99.1 | 595 |
| 3 | 14 | Lung | cfDNA | MiSeq | 11378163 | 99.1 | 330 |
| 3 | 15 | Lung | FFPE | HiSeq | 12833416 | 99.85 | 640 |
| 3 | 15 | Lung | cfDNA | HiSeq | 48343010 | 99.92 | 512 |
| 3 | 16 | Lung | FFPE | MiSeq | 8038419 | 99.32 | 691 |
| 3 | 16 | Lung | cfDNA | MiSeq | 2973720 | 99.46 | 364 |
| 3 | 17 | Lung | FFPE | HiSeq | 11109068 | 99.93 | 591 |
| 3 | 17 | Lung | cfDNA | HiSeq | 59335783 | 99.96 | 977 |
| 3 | 18 | Lung | FFPE | HiSeq + MiSeq | 83910995 | 99.49 | 1461 |
| 3 | 18 | Lung | cfDNA | HiSeq + MiSeq | 79650362 | 99.53 | 808 |
| 3 | 19 | Lung | FFPE | MiSeq | 3672913 | 99.02 | 377 |
| 3 | 19 | Lung | cfDNA | MiSeq | 7792509 | 98.94 | 169 |
| 3 | 20 | Lung | FFPE | HiSeq | 12456063 | 99.94 | 504 |
| 3 | 20 | Lung | cfDNA | HiSeq | 38831522 | 99.78 | 284 |

Supplementary Table 1. Details of all sequencing for pilot studies 1, 2 and 3.

| Sample ID | Cancer Type | Sample Type | Gene | Chromosome | Start position | End position | Reference allele sequence | Alternative allele sequence | Depth | Alt Depth | AF (%) | Alternative Technology AF (%) |
|---|---|---|---|---|---|---|---|---|---|---|---|---|
| 1 | Colon | FFPE | KRAS | chr12 | 25245350 | 25245350 | C | T | 82 | 7 | 8.5 | WT |
| 1 | Colon | FFPE | NRAS | chr1 | 114713909 | 114713909 | G | T | 172 | 92 | 53.5 | 56.3 |
| 1 | Colon | FFPE | PIK3CA | chr3 | 179218303 | 179218303 | G | A | 56 | 37 | 66.1 | 70 |
| 2 | Colon | FFPE | PIK3CA | chr3 | 179218294 | 179218294 | G | A | 400 | 119 | 29.8 | 24.2 |
| 3 | Colon | FFPE | KRAS | chr12 | 25227341 | 25227341 | T | G | 1123 | 474 | 42.2 | 47.6 |
| 4 | Colon | FFPE | KRAS | chr12 | 25227342 | 25227342 | T | A | 67 | 16 | 23.9 | 42.5 |
| 4 | Colon | FFPE | PIK3CA | chr3 | 179218294 | 179218294 | G | A | 26 | 2 | 7.7 | WT |
| 5 | Colon | FFPE | KRAS | chr12 | 25245350 | 25245351 | CC | AG | 89 | 35 | 39.3 | 30.3 |
| 7 | Colon | FFPE | KRAS | chr12 | 25225628 | 25225628 | C | T | 335 | 95 | 28.4 | 34 |
| 7 | Colon | FFPE | PIK3CA | chr3 | 179234297 | 179234297 | A | G | 164 | 23 | 14.0 | 13 |
| 8 | Colon | FFPE | KRAS | chr12 | 25245347 | 25245347 | C | T | 128 | 73 | 57.0 | 93 |
| 9 | Colon | FFPE | BRAF | chr7 | 140753336 | 140753336 | A | T | 215 | 30 | 14.0 | 19.8 |
| 9 | Colon | FFPE | TP53 | chr17 | 7,670,716 | 7,670,716 | G | A | 66 | 14 | 21.0 | 16 |
| 10 | Colon | FFPE | BRAF | chr7 | 140753336 | 140753336 | A | T | 239 | 74 | 31.0 | 34.7 |
| 11 | Colon | FFPE | KRAS | chr12 | 25225628 | 25225628 | C | T | 246 | 102 | 41.5 | 34 |
| 11 | Colon | FFPE | PIK3CA | chr3 | 179234297 | 179234297 | A | G | 88 | 41 | 46.6 | 48 |
| 11 | Colon | FFPE | TP53 | chr17 | 7675119 | 7675119 | C | T | 95 | 29 | 31.0 | 30 |
| 12 | Colon | FFPE | KRAS | chr12 | 25245350 | 25245350 | C | T | 38 | 5 | 13.2 | 8.9 |
| 12 | Colon | FFPE | PIK3CA | chr3 | 179218294 | 179218294 | G | A | 27 | 2 | 7.4 | WT |
| 13 | Colon | FFPE | KRAS | chr12 | 25245350 | 25245350 | C | A | 1213 | 294 | 24.2 | 22 |
| 14 | Colon | FFPE | PIK3CA | chr3 | 179218303 | 179218303 | G | A | 306 | 158 | 51.6 | 47 |
| 14 | Colon | FFPE | TP53 | chr17 | 7675208 | 7675208 | G | A | 388 | 18 | 18.0 | 17 |

Supplementary Table 2. A list of variants from clinical samples for pilot study 1.

| Sample ID | Cancer Type | Sample Type | Gene | Chr | Start position | End position | Reference allele sequence | Alternative allele sequence | Depth | Alt Depth | AF (%) | Alternative Technology AF (%) |
|---|---|---|---|---|---|---|---|---|---|---|---|---|
| 1 | Lung | cfDNA | EGFR | chr7 | 55191822 | 55191822 | T | G | 675 | 33 | 4.9 | 2 |
| 2 | Lung | cfDNA | EGFR | chr7 | 55181378 | 55181378 | C | T | 920 | 10 | 1.1 | 47.8 |
| 2 | Lung | cfDNA | EGFR | chr7 | 55191822 | 55191822 | T | G | 1091 | 11 | 1.0 | 44.5 |
| 3 | Lung | cfDNA | EGFR | chr7 | 55174771 | 55174786 | AGGAATTAAGAGAAGC | A | 1024 | 334 | 32.6 | 35.7 |
| 3 | Lung | cfDNA | EGFR | chr7 | 55181378 | 55181378 | C | T | 1243 | 240 | 19.3 | 18.9 |
| 4 rep 1 | Lung | cfDNA | EGFR | chr7 | 55191822 | 55191822 | T | G | 1608 | 706 | 43.9 | 33.2 |
| 4 rep 1 | Lung | cfDNA | KRAS | chr12 | 25245351 | 25245351 | C | T | 2645 | 440 | 16.6 | 15.3 |
| 4 rep 2 | Lung | cfDNA | EGFR | chr7 | 55191822 | 55191822 | T | G | 1549 | 627 | 40.5 | 33.2 |
| 4 rep 2 | Lung | cfDNA | KRAS | chr12 | 25245351 | 25245351 | C | T | 2582 | 456 | 17.7 | 15.3 |
| 4 rep 3 | Lung | cfDNA | EGFR | chr7 | 55191822 | 55191822 | T | G | 1558 | 642 | 41.2 | 33.2 |
| 4 rep 3 | Lung | cfDNA | KRAS | chr12 | 25245351 | 25245351 | C | T | 2615 | 445 | 17.2 | 15.3 |
| 5 | Lung | cfDNA | EGFR | chr7 | 55174776 | 55174794 | TTAAGAGAAGCAACATCTC | T | 862 | 32 | 3.7 | 2.4 |
| 6 | Lung | cfDNA | EGFR | chr7 | 55174772 | 55174792 | GGAATTAAGAGAAGCAACATC | AAT | 1183 | 36 | 3.0 | 4.7 |
| 7 | Lung | cfDNA | EGFR | chr7 | 55174771 | 55174786 | AGGAATTAAGAGAAGC | A | 1968 | 779 | 39.6 | 19.7 |
| 9 | Lung | FFPE | EGFR | chr7 | 55174776 | 55174788 | TTAAGAGAAGCAA | C | 2453 | 830 | 33.8 | 70 |
| 10 | Lung | FFPE | EGFR | chr7 | 55174771 | 55174786 | AGGAATTAAGAGAAGC | A | 1440 | 393 | 27.3 | 33.8 |
| 11 | Lung | FFPE | EGFR | chr7 | 55191822 | 55191822 | T | G | 958 | 384 | 40.1 | 21.2 |
| 12 rep 1 | Lung | FFPE | NRAS | chr1 | 114713909 | 114713909 | G | T | 2847 | 725 | 25.0 | 20.9 |
| 12 rep 2 | Lung | FFPE | NRAS | chr1 | 114713909 | 114713909 | G | T | 2760 | 729 | 26.4 | 20.9 |
| 12 rep 3 | Lung | FFPE | NRAS | chr1 | 114713909 | 114713909 | G | T | 2625 | 669 | 24.5 | 20.9 |
| 13 | Lung | FFPE | APC | chr5 | 112838329 | 112838329 | T | A | 97 | 17 | 17.5 | 9.9 |
| 13 | Lung | FFPE | FBXW7 | chr4 | 152324294 | 152324294 | G | A | 262 | 16 | 8.1 | 7 |
| 13 | Lung | FFPE | FBXW7 | chr4 | 152326214 | 152326214 | C | T | 2957 | 242 | 6.1 | 7.8 |
| 13 | Lung | FFPE | KRAS | chr12 | 25245351 | 25245351 | C | A | 3202 | 265 | 8.3 | 7.8 |
| 14 | Lung | FFPE | EGFR | chr7 | 55191822 | 55191822 | T | G | 5727 | 1816 | 31.0 | 32 |

Supplementary Table 3. A list of variants from clinical samples for pilot study 2.

| | | | | | | | | ATOM-Seq | | | Alternative Technology | | | |
|---|---|---|---|---|---|---|---|---|---|---|---|---|---|---|
| Sample ID | Sample Type | Gene | Chr | Start position | End position | Reference allele sequence | Alternative allele sequence | Depth | Alt Depth | AF (%) | Depth | AltDepth | AF (%) | Using UMI |
| 1 | FFPE | FGFR1 | chr8 | 38414245 | 38414245 | C | G | 57 | 4 | 7.0 | Not Covered | | | |
| 1 | cfDNA | | | | | | | 57 | 0 | 0.0 | | | | |
| 2 | FFPE | ERBB2 | chr17 | 39724748 | 39724748 | T | TGGGCTCCCC | 1759 | 325 | 18.5 | 388 | 97 | 25.0 | yes |
| 2 | cfDNA | | | | | | | 2449 | 0 | 0.0 | Not Done | | | |
| 3 | FFPE | TP53 | chr17 | 7673803 | 7673803 | G | A | 149 | 116 | 77.9 | 3087 | 1613 | 52.25 | no |
| 3 | cfDNA | | | | | | | 84 | 3 | 3.6 | Not Done | | | |
| 4 | FFPE | TP53 | chr17 | 7674917 | 7674917 | T | C | 767 | 58 | 7.6 | 164 | 13 | 7.93 | yes |
| 4 | cfDNA | | | | | | | 159 | 0 | 0.0 | 237 | 0 | 0 | yes |
| 4 | FFPE | MAP2K1 | chr15 | 66436755 | 66436761 | CTGGAGA | C | 238 | 29 | 12.2 | 458 | 30 | 6.5 | |
| 4 | cfDNA | | | | | | | 54 | 2 | 3.7 | 232 | 0 | 0 | |
| 5 | FFPE | TP53 | chr17 | 7675122 | 7675122 | T | A | 785 | 246 | 31.3 | 3819 | 1316 | 34.46 | no |
| 5 | cfDNA | | | | | | | 877 | 5 | 0.6 | Not Done | | | |
| 5 | FFPE | KEAP1 | chr19 | 10491620 | 10491620 | C | A | 1015 | 389 | 38.3 | Not Covered | | | |
| 5 | cfDNA | | | | | | | 715 | 3 | 0.4 | | | | |
| 6 | FFPE | RB1 | chr13 | 48459770 | 48459770 | G | A | 117 | 72 | 61.5 | Not Covered | | | |
| 6 | cfDNA | | | | | | | 120 | 8 | 6.7 | | | | |
| 6 | FFPE | TP53 | chr17 | 7674885 | 7674885 | C | T | 301 | 205 | 68.1 | 5591 | 4115 | 73.6 | no |
| 6 | cfDNA | | | | | | | 645 | 72 | 11.2 | Not Done | | | |
| 7 | FFPE | TP53 | chr17 | 7674872 | 7674872 | T | C | 428 | 116 | 27.1 | 4098 | 1265 | 30.87 | no |
| 7 | cfDNA | | | | | | | 553 | 5 | 0.9 | Not Done | | | |
| 7 | FFPE | TP53 | chr17 | 7675122 | 7675122 | T | C | 1052 | 438 | 41.6 | 233 | 69 | 29.61 | no |
| 7 | cfDNA | | | | | | | 1149 | 33 | 2.9 | Not Done | | | |
| 8 | FFPE | ERBB3 | chr12 | 56088574 | 56088574 | C | T | 300 | 54 | 18.0 | Not Covered | | | |
| 8 | cfDNA | | | | | | | 202 | 0 | 0.0 | | | | |
| 8 | FFPE | TP53 | chr17 | 7675135 | 7675139 | GGCGC | G | 816 | 269 | 33.0 | 3563 | 1362 | 38.2 | no |

| | | | | | | | | | | | | | | |
|---|---|---|---|---|---|---|---|---|---|---|---|---|---|---|
| 8 | cfDNA | | | | | | | 715 | 0 | 0.0 | Not Done | | | |
| 9 | FFPE | MSH2 | chr2 | 47410292 | 47410292 | G | T | 329 | 21 | 6.4 | 460 | 92 | 20 | |
| 9 | cfDNA | | | | | | | 280 | 3 | 1.1 | 67 | 3 | 4.48 | |
| 9 | FFPE | KEAP1 | chr19 | 10491944 | 10491944 | G | A | 126 | 19 | 15.1 | 280 | 69 | 24.45 | |
| 9 | cfDNA | | | | | | | 114 | 1 | 0.9 | 44 | 0 | 0 | |
| 9 | FFPE | TP53 | chr17 | 7673802 | 7673802 | C | T | 591 | 178 | 30.1 | 280 | 68 | 24.29 | yes |
| 9 | cfDNA | | | | | | | 277 | 6 | 2.2 | 68 | 0 | 0 | yes |
| 10 | FFPE | TP53 | chr17 | 7675088 | 7675088 | C | T | 789 | 68 | 8.6 | 7234 | 653 | 9.0 | no |
| 10 | cfDNA | | | | | | | 272 | 0 | 0.0 | Not Done | | | |
| 10 | FFPE | TP53 | chr17 | 7674241 | 7674241 | G | C | 808 | 89 | 11.0 | 13085 | 1346 | 10.3 | no |
| 10 | cfDNA | | | | | | | 3044 | 6 | 0.2 | Not Done | | | |
| 10 | FFPE | PIK3CA | chr3 | 179218307 | 179218307 | A | C | 1707 | 519 | 30.4 | 7274 | 2142 | 29.44 | no |
| 10 | cfDNA | | | | | | | 440 | 7 | 1.6 | Not Done | | | |
| 11 | FFPE | TP53 | chr17 | 7674247 | 7674247 | T | C | 607 | 66 | 10.9 | Not Covered | | | |
| 11 | cfDNA | | | | | | | 2767 | 0 | 0.0 | | | | |
| 11 | FFPE | JAK3 | chr19 | 17834899 | 17834899 | C | G | 125 | 17 | 13.6 | Not Covered | | | |
| 11 | cfDNA | | | | | | | 155 | 14 | 9.0 | | | | |
| 11 | FFPE | BRCA1 | chr17 | 43093010 | 43093010 | G | A | 114 | 25 | 21.9 | Not Covered | | | |
| 11 | cfDNA | | | | | | | 287 | 109 | 38.0 | | | | |
| 12 | FFPE | TSC2 | chr16 | 2064406 | 2064406 | C | T | 342 | 35 | 10.2 | Not Covered | | | |
| 12 | cfDNA | | | | | | | 1293 | 0 | 0.0 | | | | |
| 12 | FFPE | RB1 | chr13 | 48465204 | 48465205 | GC | G | 263 | 64 | 24.3 | Not Covered | | | |
| 12 | cfDNA | | | | | | | 27 | 2 | 7.4 | | | | |
| 12 | FFPE | TP53 | chr17 | 7676381 | 7676381 | C | A | 184 | 74 | 40.2 | 8559 | 3320 | 38.78 | no |
| 12 | cfDNA | | | | | | | 190 | 1 | 0.5 | Not Done | | | |
| 13 | FFPE | TP53 | chr17 | 7675994 | 7675994 | C | A | 711 | 297 | 41.8 | 1817 | 851 | 46.86 | no |
| 13 | cfDNA | | | | | | | 264 | 5 | 1.9 | Not Done | | | |
| 13 | FFPE | BRAF | chr7 | 140749350 | 140749350 | T | C | 536 | 229 | 42.7 | Not Covered | | | |
| 13 | cfDNA | | | | | | | 110 | 0 | 0.0 | | | | |

| | | | | | | | | | | | | | | |
|---|---|---|---|---|---|---|---|---|---|---|---|---|---|---|
| 13 | FFPE | KEAP1 | chr19 | 10499551 | 10499551 | C | A | 150 | 65 | 43.3 | Not Covered | | | |
| 13 | cfDNA | | | | | | | 0 | 0 | 0.0 | | | | |
| 14 | FFPE | CTNNB1 | chr3 | 41235749 | 41235749 | T | G | 871 | 75 | 8.6 | Not Covered | | | |
| 14 | cfDNA | | | | | | | 492 | 3 | 0.6 | | | | |
| 14 | FFPE | TP53 | chr17 | 7670694 | 7670694 | C | A | 652 | 164 | 25.2 | 824 | 164 | 19.9 | |
| 14 | cfDNA | | | | | | | 580 | 8 | 1.4 | Not Done | | | |
| 14 | FFPE | DMD1 | chrX | 32501785 | 32501785 | C | A | 330 | 85 | 25.8 | Not Covered | | | |
| 14 | cfDNA | | | | | | | 104 | 0 | 0.0 | | | | |
| 15 | FFPE | IDH1 | chr2 | 208248468 | 208248468 | G | A | 890 | 143 | 16.1 | Not Covered | | | |
| 15 | cfDNA | | | | | | | 241 | 58 | 24.1 | | | | |
| 15 | FFPE | KEAP1 | chr19 | 10491954 | 10491954 | C | T | 57 | 25 | 43.9 | Not Covered | | | |
| 15 | cfDNA | | | | | | | 247 | 4 | 1.6 | | | | |
| 15 | FFPE | KEAP1 | chr19 | 10489702 | 10489702 | C | G | 90 | 70 | 77.8 | Not Covered | | | |
| 15 | cfDNA | | | | | | | 190 | 13 | 6.8 | | | | |
| 15 | FFPE | TP53 | chr17 | 7675139 | 7675139 | C | A | 1005 | 838 | 83.4 | 2901 | 2517 | 86.76 | no |
| 15 | cfDNA | | | | | | | 1705 | 120 | 7.0 | 0 | 0 | n/a | |
| 16 | FFPE | ATM | chr11 | 108299779 | 108299779 | A | C | 871 | 144 | 16.5 | Not Covered | | | |
| 16 | cfDNA | | | | | | | 751 | 369 | 49.1 | | | | |
| 16 | FFPE | MSH6 | chr2 | 47803442 | 47803442 | C | T | 3018 | 1055 | 35.0 | 7231 | 2326 | 32.2 | no |
| 16 | cfDNA | | | | | | | 2524 | 8 | 0.3 | Not Done | | | |
| 16 | FFPE | BRCA2 | chr13 | 32337717 | 32337745 | CAGGAAGTCAGTTTG AATTTACTCAGTTT | C | 2045 | 817 | 40.0 | Not Covered | | | |
| 16 | cfDNA | | | | | | | 406 | 2 | 0.5 | | | | |
| 16 | FFPE | TP53 | chr17 | 7675076 | 7675076 | T | A | 743 | 326 | 43.9 | 7641 | 4178 | 54.68 | no |
| 16 | cfDNA | | | | | | | 775 | 1 | 0.1 | Not Done | | | |
| 17 | FFPE | MAP2K2 | chr19 | 4101064 | 4101064 | G | T | 120 | 16 | 13.3 | Not Covered | | | |
| 17 | cfDNA | | | | | | | 357 | 17 | 4.8 | | | | |
| 17 | FFPE | ROS1 | chr6 | 117317224 | 117317224 | C | A | 341 | 58 | 17.0 | 1910 | 339 | 17.7 | yes |
| 17 | cfDNA | | | | | | | 195 | 0 | 0.0 | 183 | 0 | 0 | yes |
| 17 | FFPE | CDKN2A | chr9 | 21971187 | 21971187 | G | A | 900 | 231 | 25.7 | 1020 | 433 | 42.5 | yes |

| | | | | | | | | | | | | | | |
|---|---|---|---|---|---|---|---|---|---|---|---|---|---|---|
| 17 | cfDNA | | | | | | | 505 | 2 | 0.4 | 234 | 0 | 0 | yes |
| 17 | FFPE | SMARC4 | chr19 | 11033317 | 11033317 | C | T | 667 | 180 | 27.0 | 698 | 335 | 48.0 | yes |
| 17 | cfDNA | | | | | | | 1261 | 15 | 1.2 | 118 | 2 | 1.69 | yes |
| 17 | FFPE | TP53 | chr17 | 7675085 | 7675085 | C | A | 2405 | 694 | 28.9 | 308 | 117 | 38.0 | yes |
| 17 | cfDNA | | | | | | | 2112 | 11 | 0.5 | 176 | 0 | 0 | yes |
| 18 | FFPE | MAP2K2 | chr19 | 4117520 | 4117520 | T | C | 462 | 55 | 11.90 | Not Covered | | | |
| 18 | cfDNA | | | | | | | 1422 | 0 | 0.0 | | | | |
| 18 | FFPE | CSF1R | chr5 | 150054182 | 150054203 | CGCTGCCACCG CTTCTGCTGCT | C | 22 | 4 | 18.2 | Not Covered | | | |
| 18 | cfDNA | | | | | | | 50 | 11 | 22.0 | | | | |
| 18 | FFPE | KEAP1 | chr19 | 10491944 | 10491944 | G | A | 119 | 26 | 21.8 | Not Covered | | | |
| 18 | cfDNA | | | | | | | 806 | 25 | 3.1 | | | | |
| 18 | FFPE | TP53 | chr17 | 7674216 | 7674216 | C | G | 1304 | 315 | 24.2 | 6192 | 2023 | 32.67 | no |
| 18 | cfDNA | | | | | | | 3269 | 74 | 2.3 | Not Done | | | |
| 18 | FFPE | STK11 | chr19 | 1220494 | 1220494 | G | T | 400 | 121 | 30.3 | Not Covered | | | |
| 18 | cfDNA | | | | | | | 633 | 17 | 2.7 | | | | |

Supplementary Table 4. A list of variants from clinical samples for pilot study 3 using paired lung cancer samples.

| Pan Caner Panel Targets | Colorectal and Lung Panel Targets | Custom Fusion Panel Targets |
|---|---|---|
| ABL1 | AMER1 | ABL |
| AKT1 | APC | ALK |
| ALK | ARAF | BRAF |
| AMER1 | BRAF | EGFR |
| APC | CDKN2A | ERBB2 |
| AR | DMD | ERG |
| ARAF | EGFR | ETV6 |
| ARID1A | EP300 | EWSR1 |
| ATM | ERBB3 | FGFR1 |
| BRAF | FBXW7 | FGFR2 |
| BRCA1 | GNAS | FGFR3 |
| BRCA2 | HRAS | FGR |
| CASP8 | KEAP1 | JAK1 |
| CCND1 | KRAS | JAK2 |
| CCND2 | MAP2K1 | JAK3 |
| CCND3 | MAP2K2 | NTRK1 |
| CDH1 | NRAS | NTRK3 |
| CDK4 | PDGFA | PDGFRB |
| CDK6 | PIK3CA | RAF |
| CDKN2A | SMAD4 | RARA |
| CHEK2 | STK11 | RET |
| CSF1R | TCF7L2 | ROS1 |
| CTNNB1 | TP53 | |
| DDR2 | | |
| DMD | | |
| EGFR | | |
| EP300 | | |
| ERBB2 | | |
| ERBB3 | | |
| ERBB4 | | |
| ESR1 | | |
| EZH2 | | |
| FBXW7 | | |
| FGFR1 | | |
| FGFR2 | | |
| FGFR3 | | |
| FGFR4 | | |
| FLT3 | | |
| GATA3 | | |
| GNA11 | | |
| GNAQ | | |
| GNAS | | |
| HNF1A | | |

| |
|---|
| HRAS |
| IDH1 |
| IDH2 |
| JAK2 |
| JAK3 |
| KDM6A |
| KDR |
| KEAP1 |
| KIT |
| KLF5 |
| KRAS |
| MAP2K1 |
| MAP2K2 |
| MET |
| MGA |
| MLH1 |
| MPL |
| MSH2 |
| MSH6 |
| MTOR |
| MYC |
| NF1 |
| NFE2L2 |
| NOTCH1 |
| NPM1 |
| NRAS |
| NTRK1 |
| NTRK3 |
| PDGFRA |
| PIK3CA |
| PTCH1 |
| PTEN |
| PTPN11 |
| RAF1 |
| RB1 |
| RBM10 |
| RET |
| RHOA |
| RIT1 |
| RNF43 |
| ROS1 |
| SETD2 |
| SF3B1 |
| SMAD2 |

| |
|---|
| SMAD4 |
| SMARCA4 |
| SMARCB1 |
| SMO |
| SRC |
| STK11 |
| TCF7L2 |
| TP53 |
| TSC1 |
| TSC2 |
| UA2F1 |
| VHL |
| ZFP36L2 |

Supplementary Table 5. A list of target genes present in all primer panels.
